# Performance comparison of Accucopy, Sequenza, and ControlFreeC

**DOI:** 10.1101/2023.09.03.556123

**Authors:** Xinping Fan, Yu S. Huang

**Affiliations:** Drug Discovery and Design Center, State Key Laboratory of Drug Research, Shanghai Institute of Materia Medica, Chinese Academy of Sciences, Shanghai 201203, China; University of Chinese Academy of Sciences, Beijing 100049, China; Genecast Biotechnology Co., Ltd., Wuxi 214105, Jiangsu, China

**Keywords:** CNA/copy number alteration, CNV/copy number variant, algorithm comparison, WGS/whole genome sequencing, cancer, tumor

## Abstract

Copy number alterations (CNAs) are an important type of genomic aberrations. It plays an important role in tumor pathogenesis and progression of cancer. It is important to detect regions of the cancer genome where copy number changes occur, which may provide clues that drive cancer progression. Deep sequencing technology provides genomic data at single-nucleotide resolution and is considered a better technique for detecting CNAs. There are currently many CNA-detection algorithms developed for whole genome sequencing (WGS) data. However, their detection capabilities have not been systematically investigated. Therefore, we selected three algorithms: **Accucopy, Sequenza**, and **ControlFreeC**, and applied them to data simulated under different settings. The results indicate that: 1) the correct inference of tumor sample purity is crucial to the inference of CNAs. If the tumor purity is wrongly inferred, the CNA detection will fail. 2) Higher sequencing depth and abundance of CNAs can improve performance. 3) Under the settings tested (sequencing depth at 5X or 30X, purity from 0.1 to 0.9, existence of subclones or not), Accucopy is the best-performing algorithm overall. For coverage=5X samples, ControlFreeC requires tumor purity to be above 50% to perform well. Sequenza can only perform well in high-coverage and more-CNA samples.

## 1 Introduction

### 1.1 What is Copy Number Alteration (cf. Copy Number Variant)?

Copy number alterations (CNAs) arise from deletions, insertions, or duplications resulting in chromosomal aberrations and aneuploidy, which are common genomic aberrations in cancer ^1^. It is notable that genomes of normal cells also exhibit variable copy numbers called germline copy number variants (CNVs) ^2^. In large studies of CNAs in cancer patients, it becomes necessary to accurately identify and separate CNAs and CNVs so as to prioritize candidate tumor suppressors and oncogenes ^2^. One of the best ways to separate CNAs and CNVs is to do tumor-normal pair sequencing and use the matched normal as the control to detect CNAs only. CNAs are often associated with specific cancer cell types ^3^, tumor aggressiveness, and patient prognosis ^4,5^. Cancer-related genes can be identified by detecting copy number alterations of tumor samples ^6^, and become targets of new treatments ^7–9^. Therefore, it is important to identify the loci and the copy number state of CNAs.

### 1.2 Technologies for Copy Number Alteration Detection

Over the years, there are many technologies developed to detect copy number alterations/variants in the genome. For example, comparative genomic hybridization (CGH) ^10^ generates an overview of chromosomal gains and losses throughout the whole genome of a tumor. In CGH, tumor DNA is labelled with a green fluorochrome, which is subsequently mixed (1:1) with red labelled normal DNA and hybridized to normal human metaphase preparations. The green and red labelled DNA fragments compete for hybridisation to their locus of origin on the chromosomes. The green to red fluorescence ratio measured along the chromosomal axis represents loss or gain of genetic material in the tumor at that specific locus. CGH technology has the advantages of using a small amount of DNA samples, only one hybridization is needed to detect genomic copy number changes, and it can be used to study both living cells, tissues, or archived tissues. However, the resolution of CGH is limited to alterations of approximately 5-10Mb. Smaller CNAs will evade its detection. The second technology is microarray-based comparative genomic hybridization (array CGH, aCGH) ^11^. Instead of using metaphase chromosomes, aCGH uses slides (known as microarray) arrayed with small segments of DNA (known as probes) as the targets for analysis. The tumor DNA is then labeled with a fluorescent dye of a specific color, while the normal DNA is labeled with a dye of a different color. The two genomic DNAs, tumor and normal, are then mixed together and applied to a microarray. Compared with CGH technology, it has two advantages. 1) Higher resolution and sensitivity. Because probes are orders of magnitude smaller than metaphase chromosomes, the theoretical resolution of aCGH is proportionally higher than that of traditional CGH. 2) Automation and programming. The aCGH technology adopts a microarray chip detection process, which can be automated by machines and computers, which is both time-saving and labor-saving. 3) Single nucleotide polymorphism(SNP) arrays ^12^. It detects gains and loss of genomic segments by detecting loss of heterozygosity (LOH). 4) Deep sequencing technology ^13^. Using deep sequencing technologies to detect copy number alterations is a relatively new field that has developed rapidly in recent years. Notably, higher sequencing depth can improve the accuracy and resolution of CNAs. More accurate breakpoints of copy number alterations can be obtained, and more forms of genomic alterations such as insertions and inversions that cannot be detected by chip-based methods can also be detected. Since deep sequencing technology does not require probes and can detect copy number alterations at single-base resolutions, it significantly increases the number of copy number alterations detected.

### 1.3 CNA-detection algorithms based on deep sequencing

With the development of sequencing technology and the reduction of sequencing cost, compared with other microarray-based technologies, deep sequencing provides higher precision and higher throughput for the analysis of copy number alteration ^13^. Many algorithms have been developed for the detection of copy number alteration. These algorithms can be divided into three categories based on the data used by the algorithms. The first category is to only use the depth information of the segment, such as CNAnorm, THetA, etc. ^14,15^. The second type is to only use single nucleotide variant information, such as PurityEst and PurBayes ^16,17^. The third category is to use both segment depth and single nucleotide variant information, such as PyLOH and patchwork ^18,19^. Identifiability problems exist for the first and second types of methods, since different combinations of tumor sample purity and cancer cell ploidy estimates may yield identical observations. For example, if the tumor purity estimate is 0.4 and the cancer cell ploidy estimate is 4, or if the tumor purity is 0.8 and the cancer cell ploidy is 3, both can result in the tumor sample ploidy to be 2.8 (hence identical tumor sequencing coverage data). Likewise, using only single-nucleotide variant information results in a similar identifiability problem. Therefore, these algorithms use explicit or implicit assumptions in order to solve the identifiability problem. For example, PurityEst, an algorithm that uses B-allele frequencies (BAFs), contains the unreasonable assumption that the BAFs value should always be close to 0.5. Therefore, the third type of methods, which simultaneously uses segment depth information and single nucleotide variant information, can better solve the identifiability problem. According to the results generated by the algorithm, the algorithm can be further divided into three subcategories. The first subcategory is to only estimate the total copy number of any segment. The second subcategory estimates both total copy number and the allele-specific copy number of any segment. Compared to the first subcategory, the second subcategory is able to infer the genotype. For example, if the total copy number of a gene is 3, and its candidate genotypes include AAA, AAB, ABB, and BBB, the first subcategory can only infer the copy number of the gene, while the second subcategory can also infer its genotype. A third subcategory has the additional ability to estimate whether a certain segment is subclonal. Since there are subclones in a tumor tissue due to tumor heterogeneity, the inference of subclonal segments is of great significance. However, inferring the copy number state of subclonal segments is challenging.

Although there are many algorithms designed for deep sequencing data, the impact of the sequencing depth, tumor purity, the extent of copy number alteration of cancer cells, and the number of subclones on these algorithms has not been systematically studied. Therefore, in this study, we compared three algorithms. We simulated whole-genome sequencing data under different purities, different numbers of subclones, varying extent of copy number alterations, and varying sequencing depths to evaluate these algorithms and find out their pros and cons.

## 2 Materials and methods

### 2.1 Algorithms selected

We conducted a literature search (early 2019) and selected the following algorithms: Accucopy, ControlFreeC, and Sequenza ^20–22^. ControlFreeC first estimates the copy number of a genomic segment, then performs an estimation of the tumor sample purity, and then corrects the prior copy number estimates to obtain the final copy number estimate of each segment. Both Accucopy and Sequenza use the likelihood function to first estimate the tumor purity and cancer-cell ploidy, and then derive the copy number of each segment. Unlike Sequenza, Accucopy can also tell if a segment is subclonal, using the posterior probability to calculate the probability that a segment is clonal. If the posterior probability is greater than 0.95, the segment is clonal and its copy number is an integer. If it is lower than 0.95, the segment is subclonal and its copy number is a floating-point number. The computing process of these software can be roughly divided into four steps: 1. Data preprocessing; 2. Normalization; 3. Segmentation; 4. Estimating copy number for each segment. The comparison of the three algorithms is shown in Table 1. We ran all software with default parameters or ones recommended by the authors in their corresponding papers.

**Table 1.**
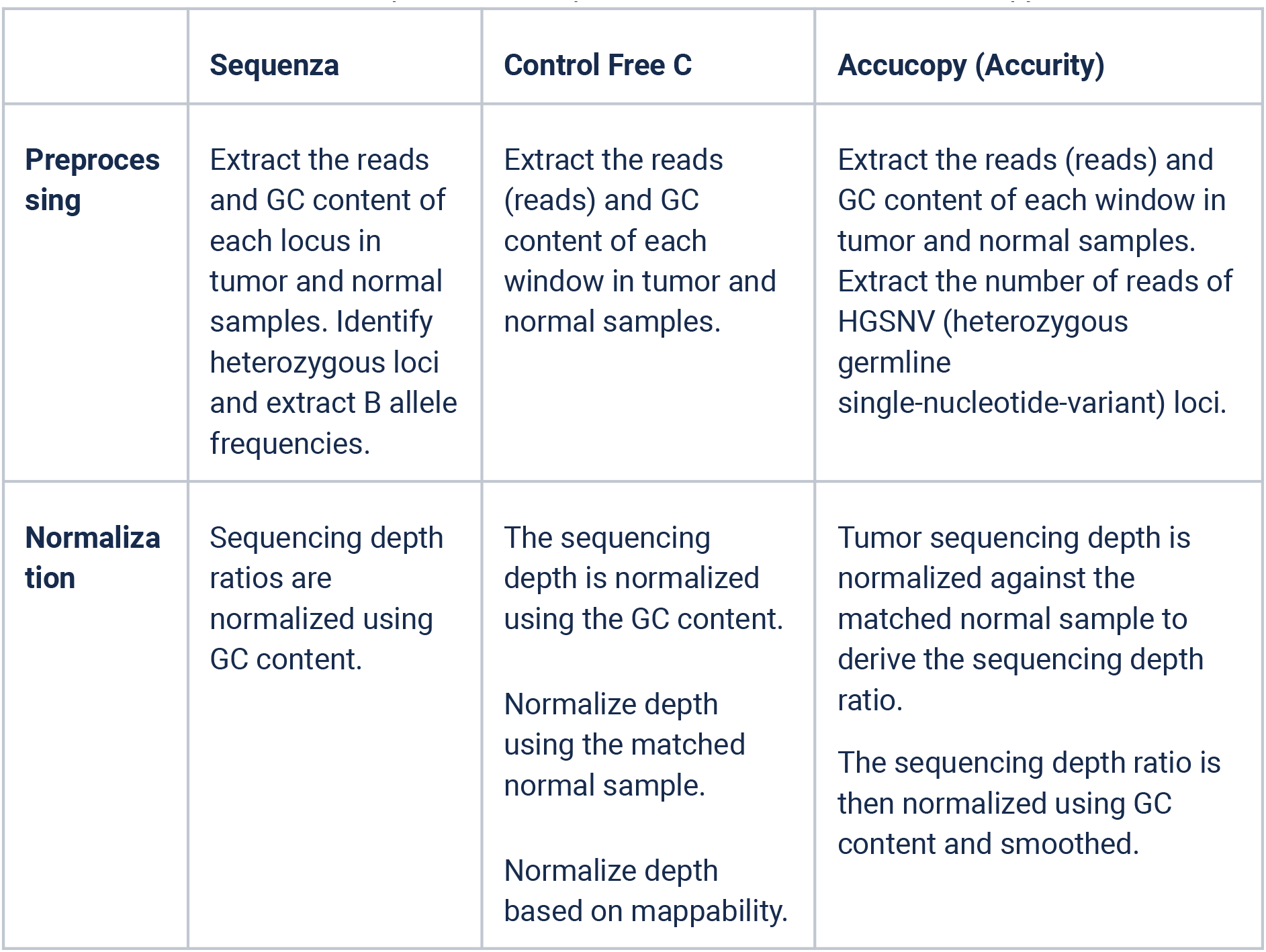
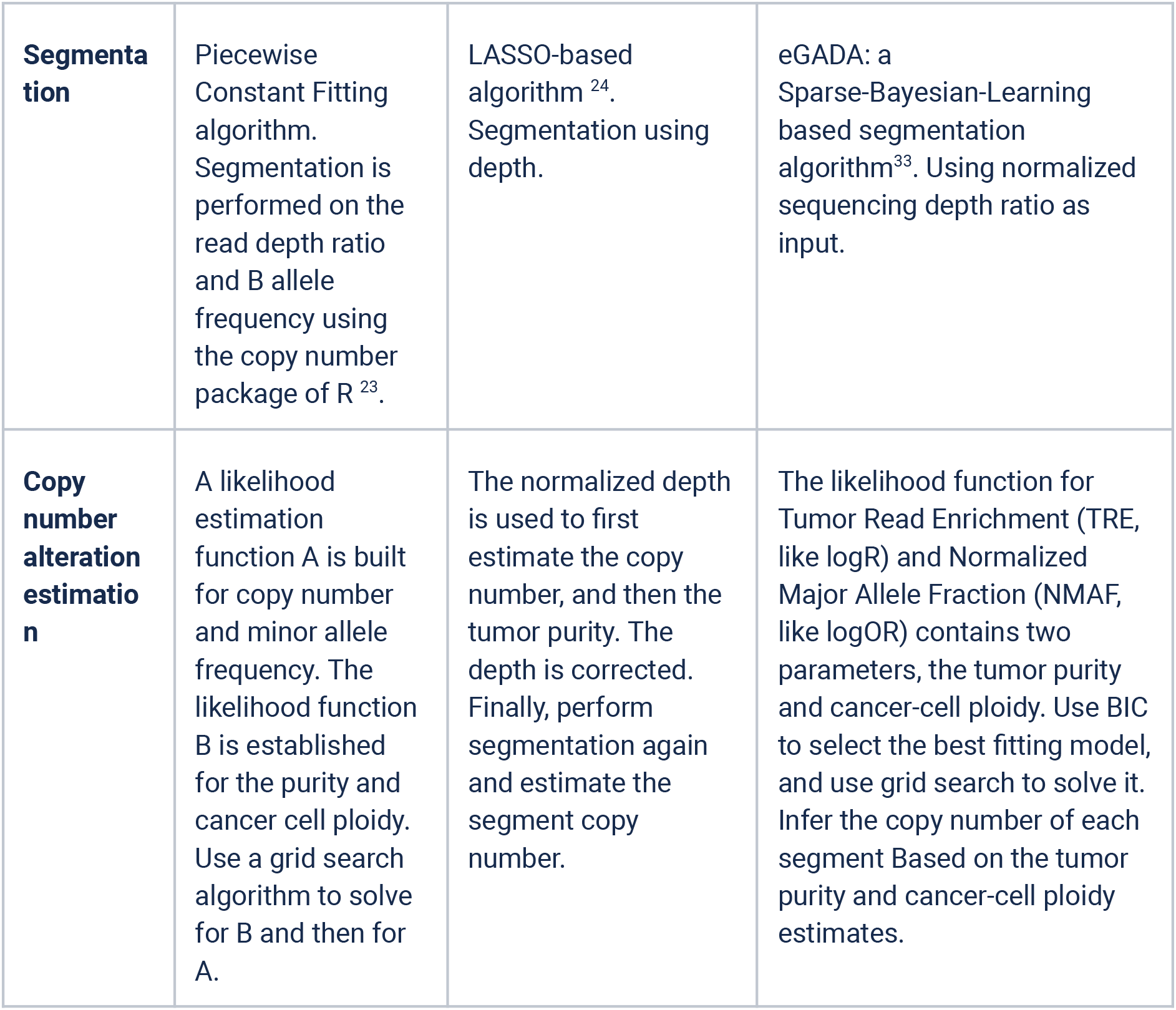
Feature comparison of Sequenza, ControlFreeC, and Accucopy.

### 2.2 Generation of simulation data

This study used EAGLE (https://github.com/sequencing/EAGLE) to simulate whole-genome sequencing data. The hg19 reference human genome is used as the reference genome for generating simulation data. EAGLE mainly uses the reference genome as a template. In addition, the simulated samples used the SNPs with MAF (major allele frequency)>10% in the 1000 Genomes Project as the single nucleotide variant (SNV) data for all samples. In this study, EAGLE is used to simulate the WGS data for both tumor and normal samples. When simulating a tumor sample, the sequencing reads of cancer cells and normal cells in the tumor sample are simulated in two steps. Copy number alterations and SNVs are introduced into the sequencing data of cancer cells, but only SNVs are introduced into normal cells. EAGLE generates the sequencing bam files for two types of cells respectively. The bam files were subsequently converted to fastq files using SamToFastq of the PICARD package. The sequencing reads of cancer cells and normal cells in the cancer tissue sample are then mixed according to a pre-set ratio to simulate a tumor sample with a non-100% purity. All the reads are aligned to the reference genome using Burrows-Wheeler Aligner (BWA) ^25^. The bam indexes are generated by samtools ^26^. Figure 1 is the flowchart for generating simulated data. In total, we generated four simulation datasets.

1. The first dataset (label: **low-cov-single-clone**) contains 9 tumor samples with sequencing coverage at 5X, no subclone, 10 CNA events, and tumor purity from 0.1 to 0.9 (step=0.1).
2. The second dataset (**low-cov-w-subclone**) contains 9 tumor samples with sequencing coverage at 5X, two or three subclones, 10 CNA events, and tumor purity from 0.1 to 0.9 (step=0.1).
3. The third dataset (**high-cov-w-subclone**) contains 3 tumor samples with sequencing coverage at 30X, tumor purity at 0.7, and containing 1, 2, or 3 subclones.
4. The fourth dataset (**low-cov–more-CNA**) contains 9 tumor samples with sequencing coverage at 5X, no subclone, tumor purity from 0.1 to 0.9 (step=0.1), a higher number of CNA events (=21).

**Figure 1.**
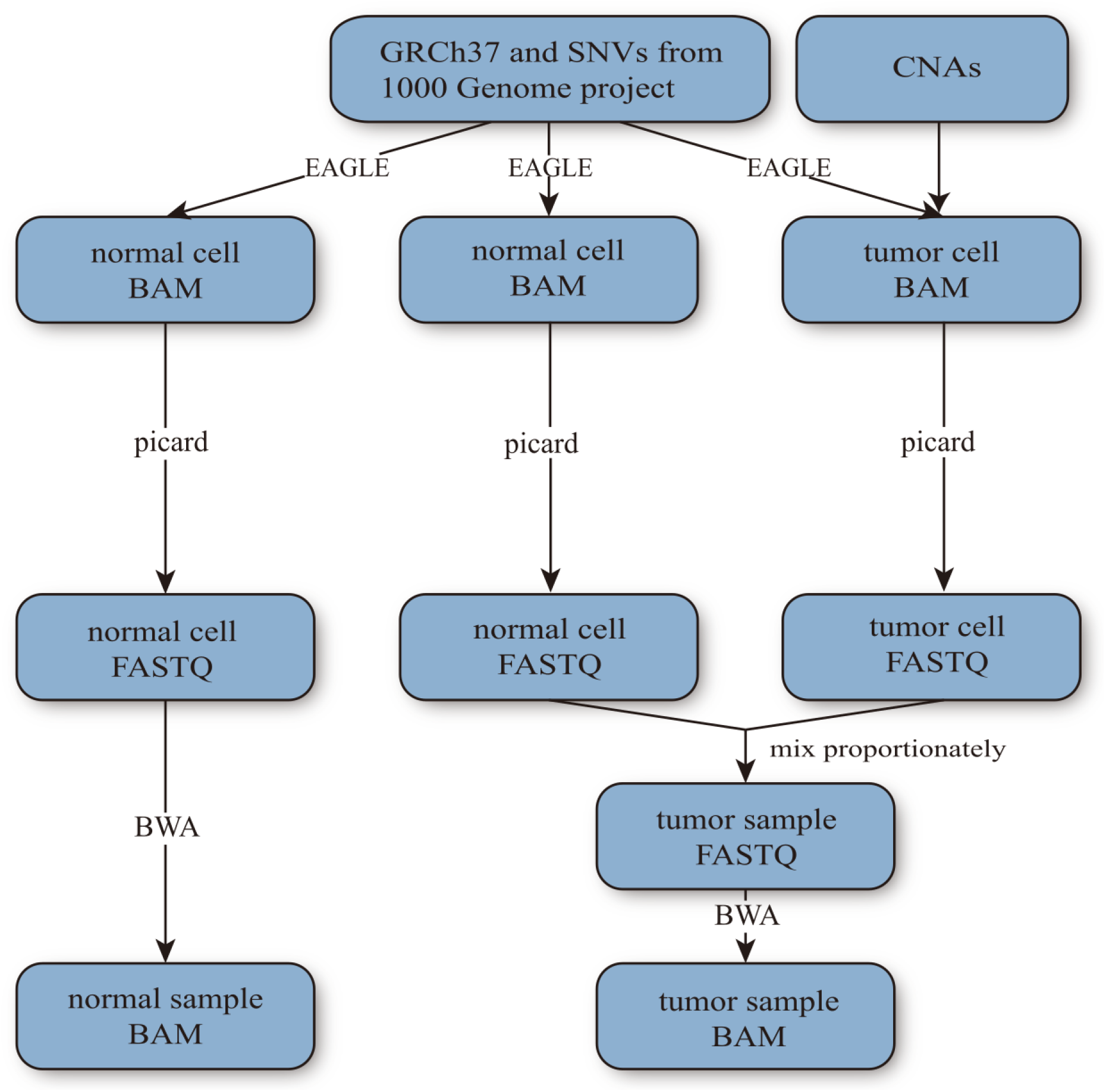
Workflow to generate the simulation data.

### 2.3 Algorithm Evaluation

We designed our evaluation metrics (precision and recall) to take both the locations of identified copy number alterations and their assigned copy number into account, while most traditional CNV-algorithm evaluation metrics only consider the locations. We use X to represent the true CNA set, and Y to represent the estimated CNA set by an algorithm. X and Y contain only non-normal (copy-number!=2) chromosomal regions. We use n and m (i and j as their indices) to denote the number of elements in set X and Y respectively. B and C represent the region (chromosome, start, end) and the copy number of a CNA respectively. Therefore:

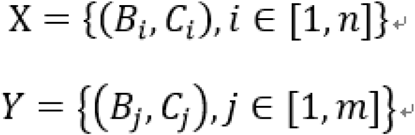

The intersection set of X and Y (X∩Y) is defined as follows. 1) If the two segments are on the same chromosome, the intersection is their overlapping portion. 2) If the two segments are not on the same chromosome, the intersection is null. The union of X and Y (X⋃Y) is similarly defined, Figure 2.

**Figure 2.**
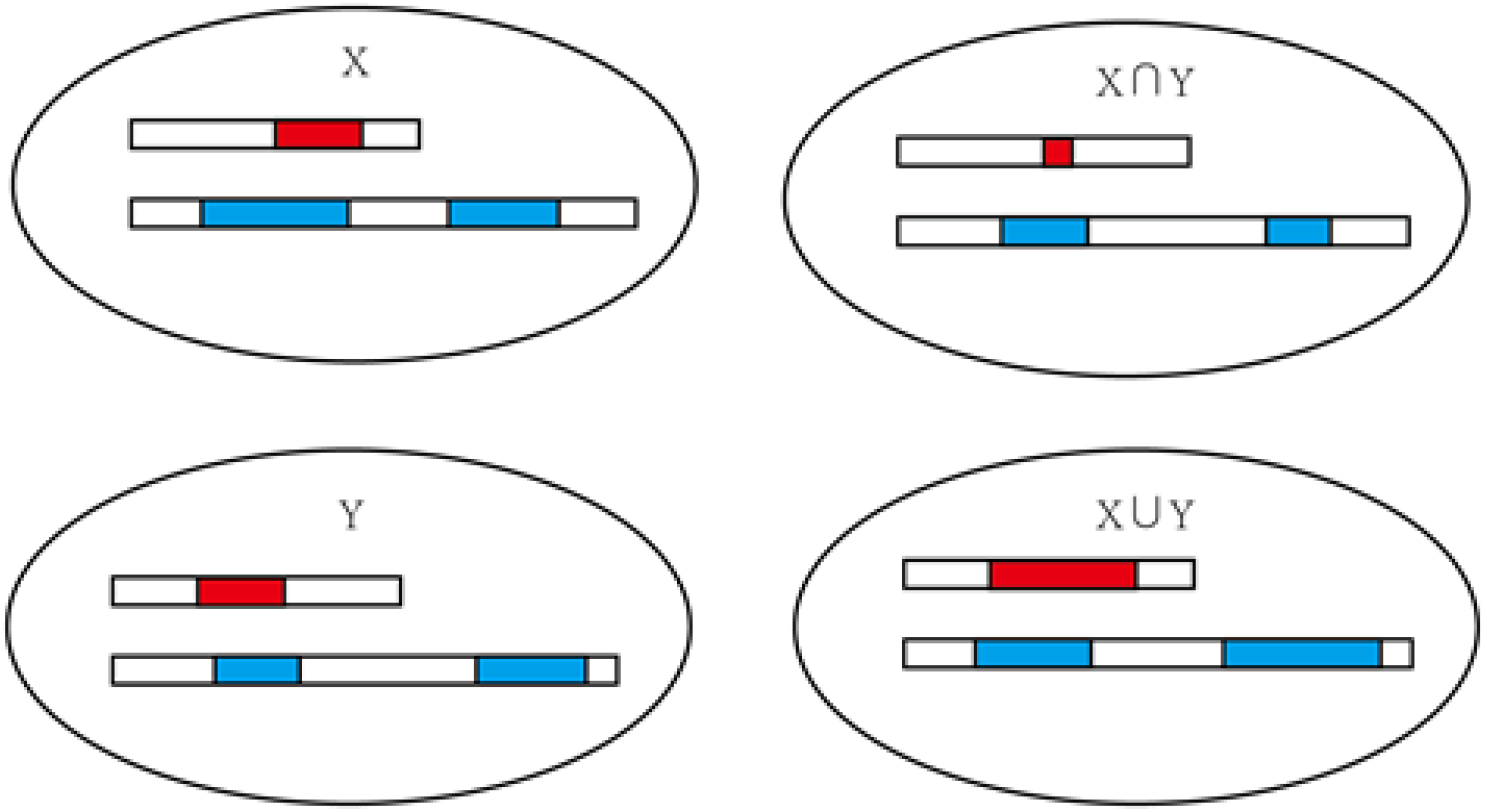
Schematic diagram of collections and their operations

The precision and recall are defined as follows. The 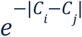 in the numerator is a metric between 0 and 1. It is 1 if two copy numbers are identical and is less than 1 if the two differ.

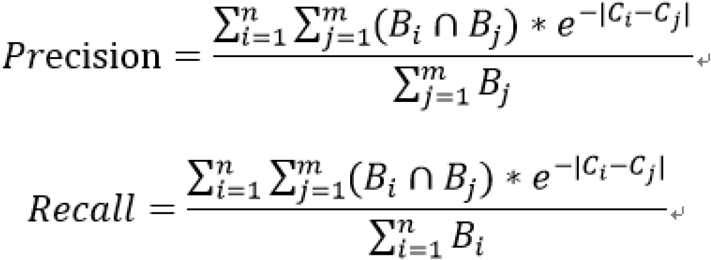

## 3 Results

### 3.1 How tumor purity impacts CNA detection accuracy

We used the first simulation dataset (low-cov-single-clone) to evaluate Accucopy, ControlFreeC, and Sequenza. Figure 3a shows the recall and precision of three algorithms, in which the Accucopy dots are concentrated in the upper right corner. The Sequenza dots are scattered, indicating that its results are unstable for this simulation data. The ControlFreeC results are concentrated either in the upper right or lower left corner, which suggests it can perform well under certain conditions but poorly under other conditions. Figure 3b plots the recall and precision of three algorithms under different tumor purity. Combining Figure a and Figure 3b, it can be concluded that the performance of Accucopy is best, with high recall and high precision regardless of the tumor purity. ControlFreeC performs well if the sample purity is above 0.6 but performs poorly if the sample purity is lower than 0.6. Sequenza seems to perform poorly at these sequencing-coverage=5X samples regardless of the tumor purity.

**Figure 3.**
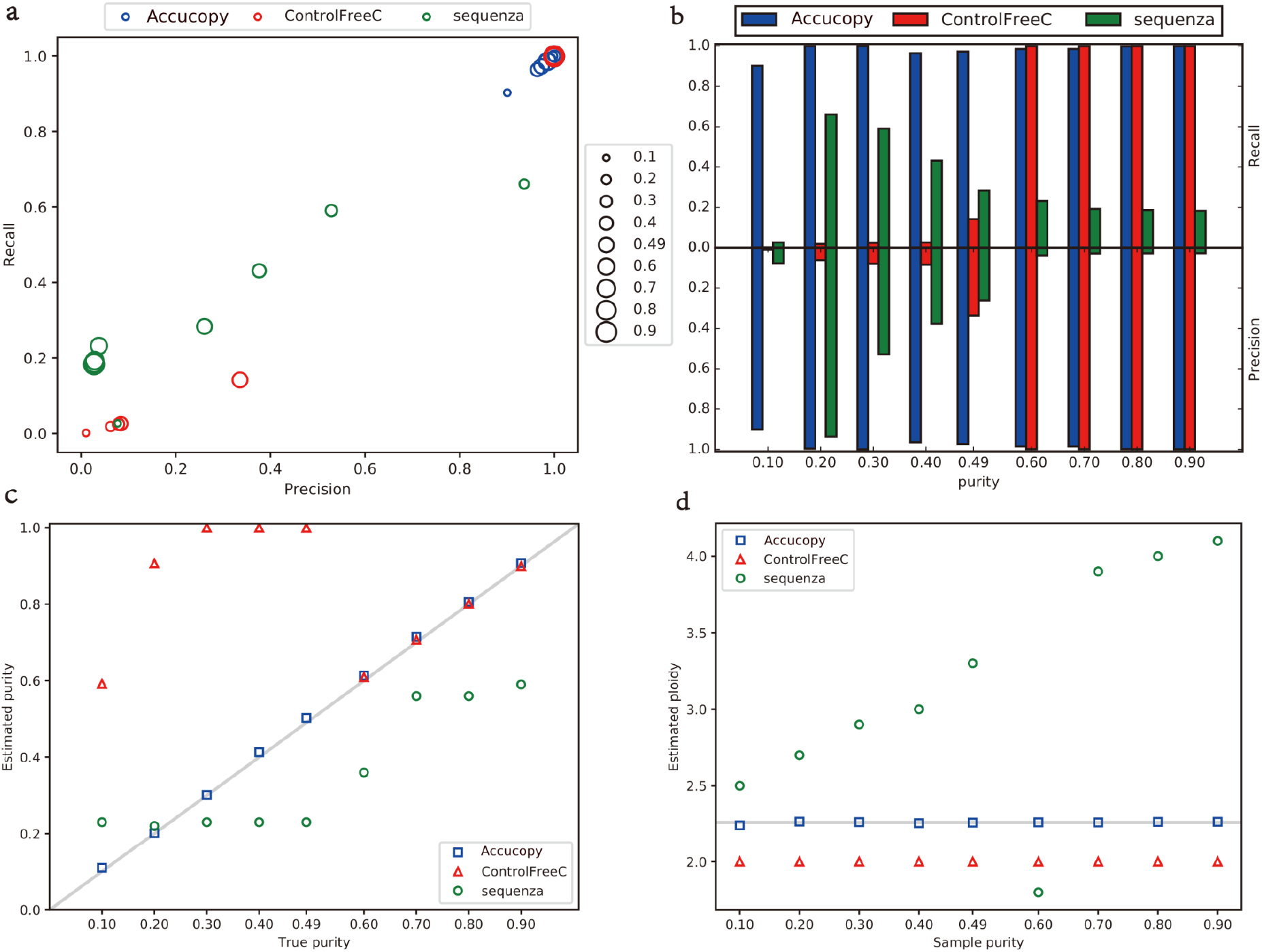
Comparison of CNA calling, tumor purity, and cancer-cell ploidy inference under different tumor purity. **a**) Scatter plots of precision and recall. Each dot represents one tumor sample. The size of each dot is proportional to its tumor purity. The colors correspond to different algorithms. **b**) Plot precision and recall as a bar plot vs true tumor purity. **c**) True tumor sample purity vs. estimated purity. **d**) True tumor purity (X-axis) vs. estimated cancer-cell ploidy. The gray line in panel d is the true cancer-cell ploidy (all simulated tumor samples are of the same cancer-cell ploidy). The closer each algorithm is to the gray line, the better it is.

Making an accurate inference about the tumor purity is critical for these algorithms. Figure 3c shows the relationship between the tumor purity estimates by three algorithms and the true value. Accucopy is the most accurate in inferring the tumor purity. The ControlFreeC estimates are only accurate if the true purity is about 0.6 or higher, which explains why its authors claim that tumor purity inference by ControlFreeC will only work if the tumor purity is higher than 50% ^21^. Figure 3d shows the relationship between the true tumor purity and the cancer-cell ploidy estimates by three algorithms. Cancer-cell ploidy estimates by Accucopy are consistently close to the true cancer-cell ploidy. ControlFreeC tends to estimate the ploidy of cancer cells always as an integer (=2). For Sequenza, its cancer-cell ploidy estimates have a strong linear relationship to tumor sample purity, the cause of which is uncertain. Considering Figure 3b and 3c together, we conclude that if an algorithm’s tumor purity estimate is close to the true tumor purity, its CNA estimates tend to be more accurate.

### 3.2 How sequencing depth impacts algorithm performance

We ran three algorithms on single-clone simulated data at sequencing depth of 5X or 30X to evaluate the impact of sequencing depth on their performance. Due to the prohibitively long running time on 30X data (esp. Sequenza), we only compared their performance on samples with tumor purity of 0.7. Table 2 shows the corresponding precision and recall of three algorithms. Figure 4a is the comparison between the inferred CNAs of three algorithms and the true CNAs at 5X, and Figure 4b is the comparison at 30X. Taken together all results, the performance of Accucopy and ControlFreeC is consistently high at 5X or 30X. However, the performance of Sequenza is pretty poor at 5X and only improves at 30X.

**Table 2.** The performance of each algorithm at different sequencing depths with true tumor purity =0.7.

| Sequencing<br>Depth | Accucopy |  | ControlFreeC |  | Sequenza |  |
| --- | --- | --- | --- | --- | --- | --- |
|  | Recall | Precision | Recall | Precision | Recall | Precision |
| 5X | 0.9855 | 0.9855 | 0.9993 | 0.9992 | 0.1929 | 0.0287 |
| 30X | 0.9638 | 0.9638 | 0.9994 | 0.9994 | 0.9535 | 0.9969 |

**Figure 4.**
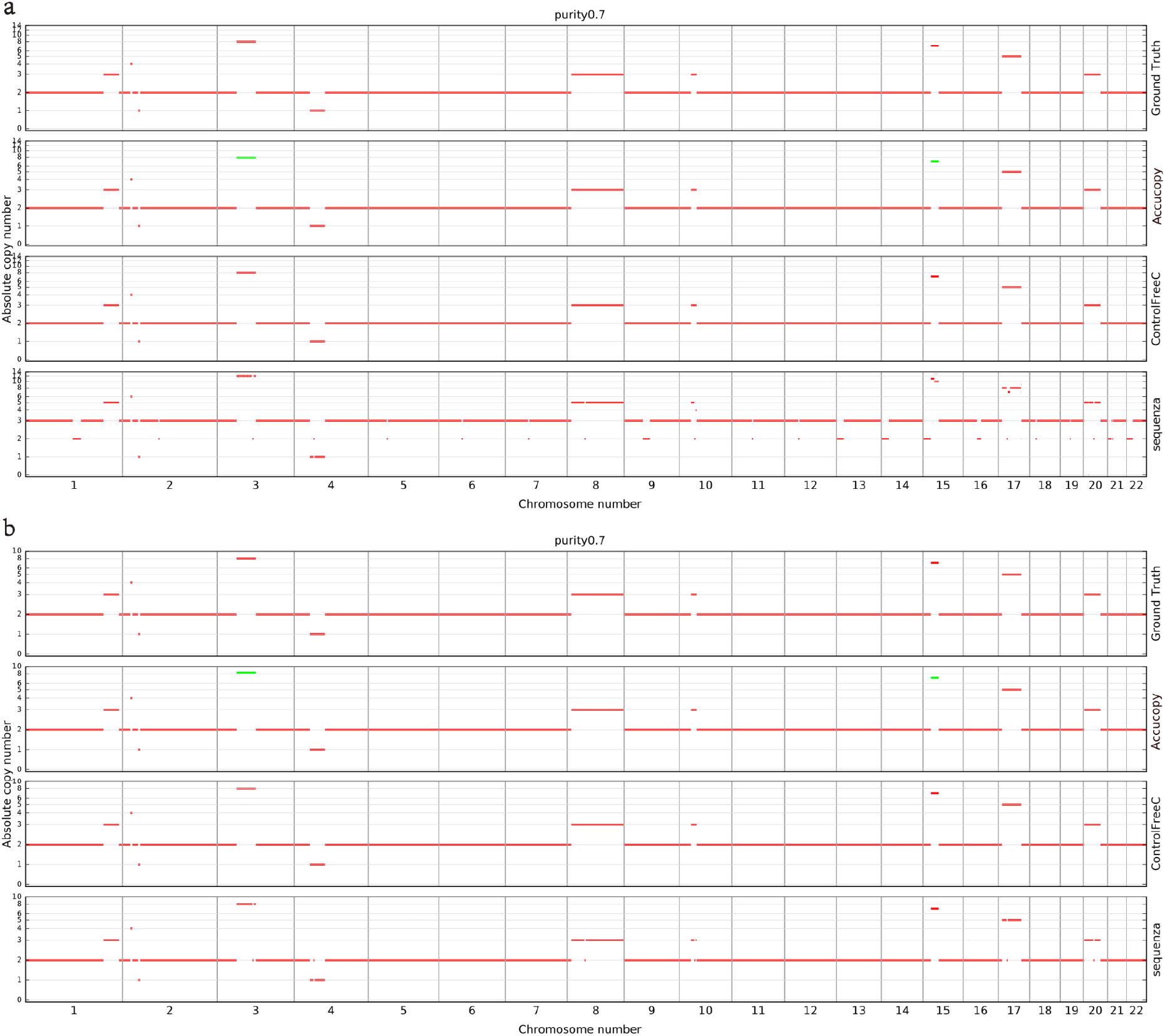
The performance of each algorithm at different sequencing depths. **a**) Sequencing depth at 5X. **b**) Sequencing depth at 30X. The CNA ground truth is plotted at the top of each panel. Red segments are clonal while the green ones are subclonal.

### 3.3 How tumor heterogeneity impacts algorithm performance

We mimicked tumor heterogeneity by simulating tumor samples with one, two, or three subclones, corresponding to the first and second simulation dataset. The comparison results are shown in Fig. 5. Fig. 5a and Fig. 5b are the results for tumor samples containing two subclones. Figure 5c and Figure 5d are the results for three subclones. The performance of Accucopy and ControlFreeC slightly decreased from two subclones to three subclones, indicated by the lower recall bar in Fig 5d vs 5b.

**Figure 5.**
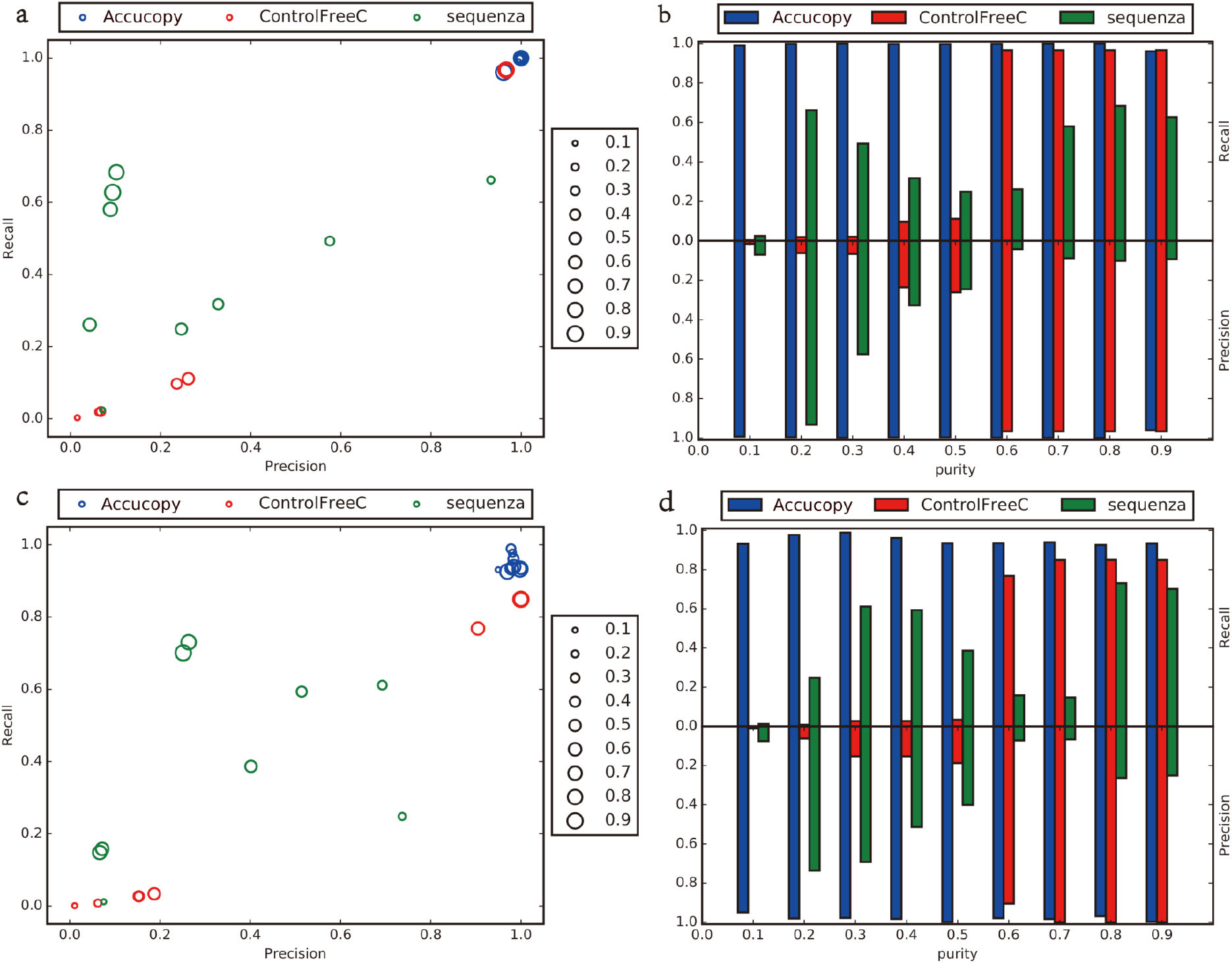
Compare performance under different subclone scenarios. **a**) Scatter plot of precision and recall of three algorithms with two subclones. The size of each dot is proportional to the purity of the corresponding sample. **b**) Barplots of precision and recall vs true tumor purity with two subclones. **c**) Scatter plot of precision and recall of three algorithms with three subclones. **b**) Barplots of precision and recall vs true tumor purity with three subclones.

Figure 6a plots the recall and precision of Accucopy, ControlFreeC, and Sequenza vs the number of subclones while the tumor purity is fixed at 0.8. Figure 6b shows a similar plot but the tumor purity is fixed at 0.9. Regarding the impact on CNA estimates, the number of subclones in a tumor sample is statistically confounded with tumor purity. For example, tumor sample 1 contains 40% normal cells and 60% monoclonal cancer cells, the copy number of locus G in normal cells is 2, the corresponding copy number in cancer cells is 4, so the observed copy number of locus G is 3.2 (=2*0.4+4*0.6). In another sample, sample 2 consists of 20% normal cells and 80% cancer cells, in which the cancer cells consist of two subclones. Cancer clone A is 40% of the entire tumor sample, and clone B is also 40%. Similarly, the copy number of locus G in normal cells is 2, its copy number in cancer clone A is 3, and its copy number in cancer clone B is 4, then the observed copy number of locus G is 3.2 (=2*0.2+3*0.4+4*0.4), the same as sample 1. But in fact, the copy number of locus G in cancer cells of sample 1 is 4, while its copy number in cancer cells of sample 2 is 3 or 4 depending on which clone.

**Figure 6.**
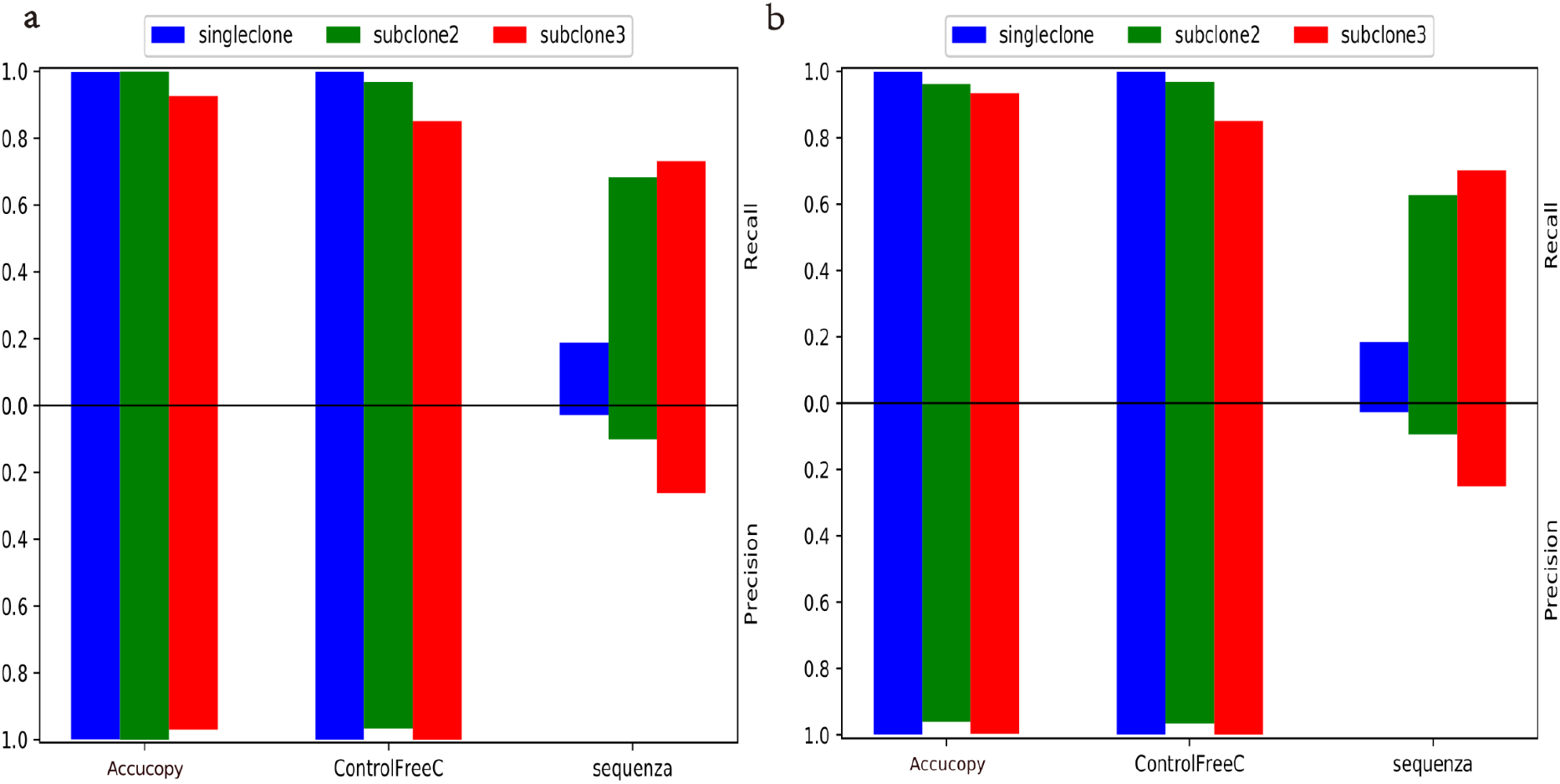
CNA recall and precision under different levels of tumor heterogeneity. **a**) Tumor sample purity=0.8. **b**) Tumor purity=0.9. The blue bar represents samples with one subclone. The green bar represents samples with two subclones. The red bar represents samples with three subclones

In general, as tumor heterogeneity (proxied by the number of subclones in a tumor sample) increases, the performance of both Accucopy and ControlFreeC decreases. Sequenza is too unreliable at low coverage (5X) to be included to draw any conclusion. Accucopy has the ability to estimate whether a segment is clonal (same CNA state across all cancer cells in a tumor sample) or subclonal, but the ability is not perfect at present. ControlFreeC does not discriminate between clonal or subclonal segments, so the number of subclones has a greater impact on ControlFreeC than Accucopy, Figure 6. Based on these results, we conclude that the performance of these algorithms decreases as the number of subclones increases.

### 3.4 How abundance of CNAs impacts algorithm performance

To investigate how abundance of CNAs impacts algorithm performance, we used two simulation datasets, low-cov-single-clone and low-cov–more-CNA. Figures 7a and 7b are the results based on the first dataset, and Figures 7c and 7d are the results based on the second dataset. Sequenza is again a bit an outlier. When tumor purity is 0.6 or 0.7, Sequenza performed significantly better with more CNAs in the sample. Sequenza performed poorly in samples with true tumor purity equal to 0.8 or 0.9 because its tumor purity and cancer-cell ploidy estimates were wrong. Overall, as the number of CNA events increases, all three algorithms improve, presumably because more CNA events can provide more information for inference.

**Figure 7.**
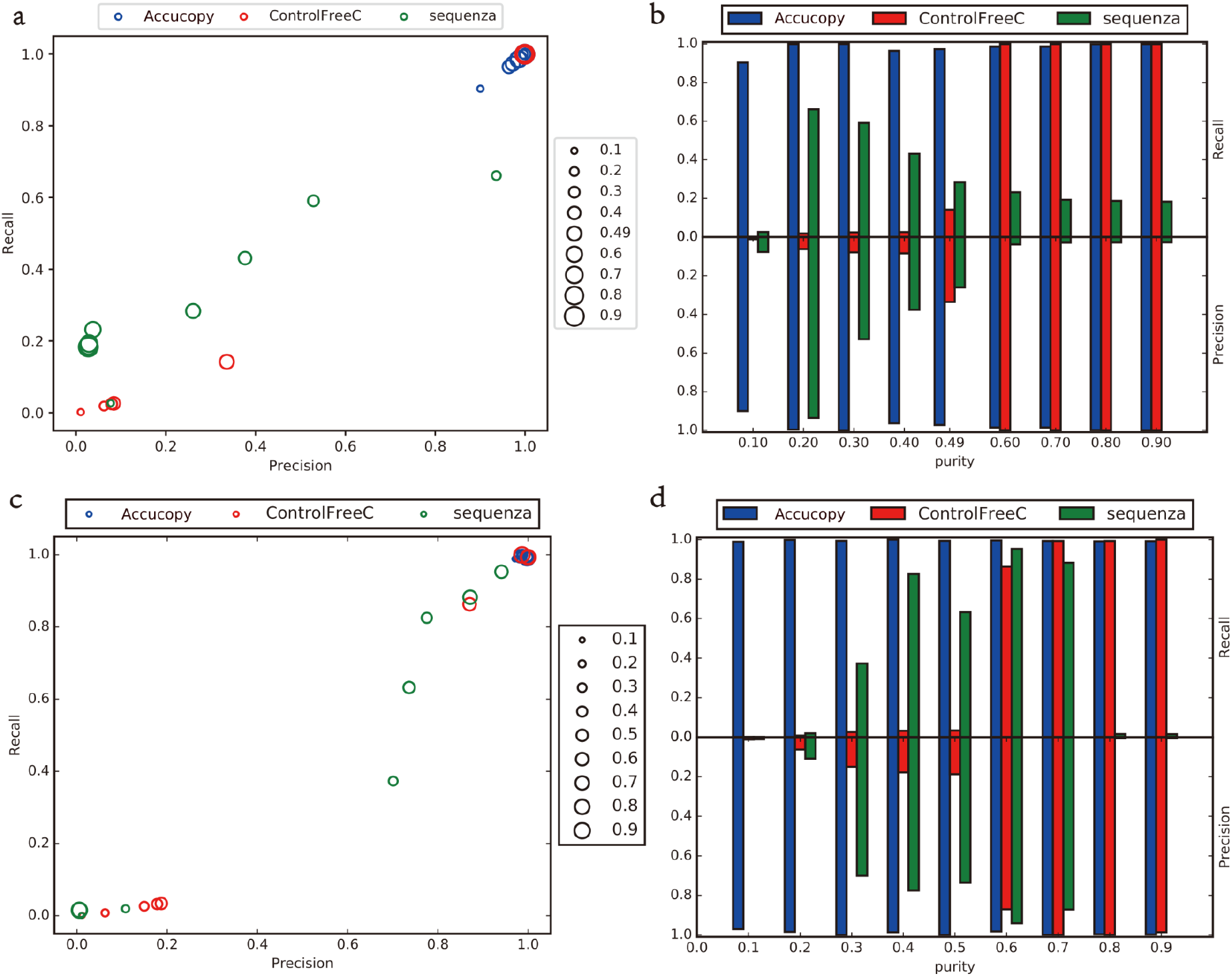
Compare algorithm performance under different numbers of CNA events. **a**) The scatter plot of precision and recall for samples with the number of CNA events equal to 10. **b**) The barplot of precision and recall with the number of CNAs equal to 10. **c**) The scatter plot of precision and recall with the number of CNA events equal to 21. **d**) The barplot of precision and recall with the number of CNAs equal to 21.

### 3.5 Running time comparison

Table 3 shows the running time of three algorithms running on 5X samples. The CPU model tested is Intel Xeon E5-2630@2.4GHz, and each program is allocated with 10GB of memory. It can be seen that the running time of Accucopy and ControlFreeC is relatively short. Both usually finish within 1-2 hours, while Sequenza takes nearly two days. This is due to not only the differences in algorithm design, but also different programming languages in which the algorithms were implemented. In terms of algorithm design, ControlFreeC does not involve a large amount of computing, while Accucopy and Sequenza have a large amount of computing due to the likelihood estimation step. Also the latter two need to search a space of parameters to find the optimal one. But Sequenza is more brute-force (exhausting the entire search space) than Accucopy (use the periodic pattern in coverage ratio to guide the search). In terms of programming languages, Accucopy and ControlFreeC were mainly written in high-performance languages such as C++ or Rust, but Sequenza was written in Python and R. Hence, Sequenza runs considerably longer than ControlFreeC or Accucopy.

**Table 3.**
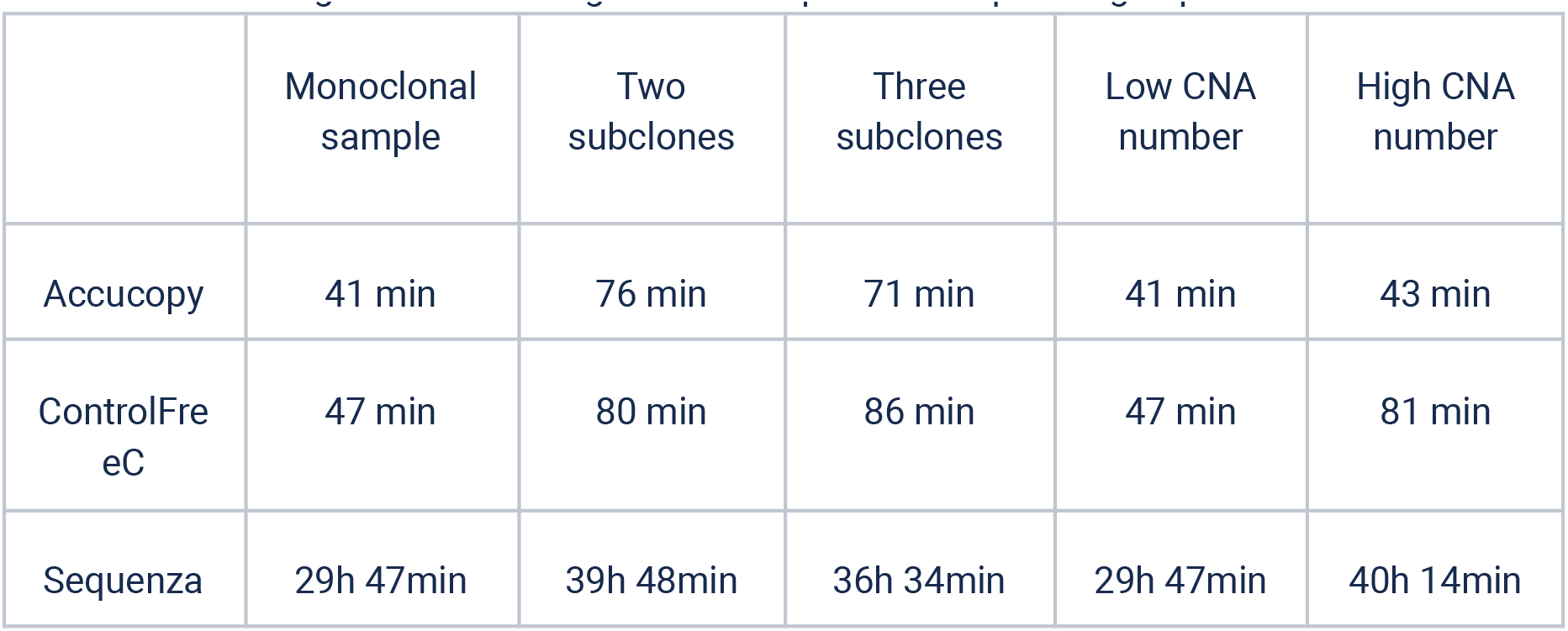
The running time of each algorithm on input with sequencing depth of 5X.

## 4 Discussion

In this study, we compared the CNA inference capability of Accucopy, ControlFreeC, and Sequenza on simulated tumor-normal WGS data. We simulated tumor samples by varying tumor purity, the number of subclones, the sequencing depth, and the number of copy number alterations in cancer cells. Based on the simulation results, Accucopy has been shown to be the best overall and has a relatively short running time. For coverage=5X samples, ControlFreeC requires the tumor purity to be above 50% to perform well. When the tumor purity is lower than 50%, its results are quite unreliable. We think this is related to the algorithm it uses to infer the tumor purity. But its running time is reasonably fast. Sequenza did not perform well in low-coverage or few-CNA settings. But if the data can provide better statistical signals, such as higher sequencing depth (i.e. 30X) and more abundant CNAs, it can produce reliable results. But it takes days to finish.

For any deep-sequencing based CNA detection algorithm, the correct inference of tumor purity is the key to its success. Besides the correctness of tumor purity estimates, users have to consider if a chosen algorithm fits your use case. For example, not all algorithms can work in low-coverage samples. Secondly, the running time of algorithms can differ by an order of magnitude. Some algorithms are slow due to complex models or the choice of slow programming languages. For three algorithms we compared: Accucopy, ControlFreeC and Sequenza. Accucopy adopts a model more complex than ControlFreeC, which is why it performs well in low-coverage and low-tumor-purity samples. Models of Accucopy and Sequenza are of similar complexity. But Accucopy is optimized to have a smaller search space, so its running time is considerably less than that of Sequenza.

The detection of subclonal segments, the inference of the genotype of subclonal segments and the inference of the number of subclones in the sample are questions that have not been thoroughly compared. Accucopy can only tell if a segment is subclonal or not, but can infer neither the number of subclones nor the genotypes in all subclones, while ControlFreeC and Sequenza do not consider subclones. Many algorithms ^27–32^ have been proposed to tackle the problem of inferring the number of subclones and genotypes of subclonal segments.

Our simulation-based comparative study is by no means comprehensive. For example, we only compared sequencing depth at 5X and 30X. The simulated samples are simple relative to clinical samples while the latter display all sorts of tumor heterogeneity and much more complex CNAs. However, we hope this limited simulation study can offer some guidance to users at large.

